# Mapping Cell Membrane Organization and Dynamics Using Soft Nano-Imprint Lithography

**DOI:** 10.1101/767590

**Authors:** T. Sansen, D. Sanchez-Fuentes, R. Rathar, A. Colom-Diego, F. El Alaoui, J. Viaud, M. Macchione, S. de Rossi, S. Matile, R. Gaudin, V. Bäcker, A. Carretero-Genevrier, L. Picas

## Abstract

Membrane shape is a key feature of many cellular processes, including cell differentiation, division, migration, and trafficking. The development of nanostructured surfaces allowing for the *in situ* manipulation of membranes in living cells is crucial to understand these processes, but this requires complicated and limited-access technologies. Here, we investigate the self-organization of cellular membranes by using a customizable and bench top method allowing to engineer 1D SiO_2_ nanopillar arrays of defined sizes and shapes on high-performance glass compatible with advanced microscopies. As a result of this original combination, we provide a mapping of the morphology-induced modulation of the cell membrane mechanics, dynamics and steady-state organization of key protein complexes implicated in cellular trafficking and signal transduction.

## Introduction

The plasma membrane constitutes a complex biochemical and mechanical structure mediating the first interaction of cells with the external environment. A key feature that determines this physiological function relies on the ability of the cell membrane to undergo changes in shape to support the interaction of cells with the external environment^1–4^. Indeed, changes in membrane morphologies, from the nanometer to the micrometer scale, take place during all the endo-or exocytic pathways but also, during other relevant cellular processes such as migration, cell-cell fusion or invasion among others, which often involve the formation of large membrane protrusions. Establishing how the plasma membrane morphology triggers biochemical signals through the recruitment, reorganization and/or disassembly of membrane-associated proteins is decisive to understand these cellular functions. A recognized strategy to investigate the cellular response to membrane topology consists in manipulating the membrane shape *in situ*. Thus, important mechanistic insight has been gathered from the technical flexibility of minimal reconstituted systems to recapitulate any organelle morphology and to quantitatively assess biochemical and mechanical responses^5,6^. Yet, the approaches allowing for equivalent biophysical studies in living cells are still limited. While microtextured surfaces made of soft polymeric materials, such as PDMS or SU-8, in the form of vertical micro-to nano-pillar arrays have been largely applied to investigate the transduction of mechanical stimuli on cells^7–9^, it is unclear whether the traction forces exerted by cells on soft pillars might induce controlled plasma membrane curvatures^10^. Also, nanostructured polymeric materials typically reproduce the physiological rigidity of the extracellular milieu^11^ although they can compromise the quality of imaging across nanopillar surfaces due to refractive index mismatching^12^. Controlled bending of cellular membranes can be achieved with materials such as SiO_2_ (Young’s modulus ∼ 64 GPa), which display optical properties compatible with advanced microscopy experiments. Whereas dispersed SiO_2_ or polystyrene beads of well-defined diameters have become a widely used strategy in biology^13–15^, they often exhibit a challenging handling on surfaces and display limited morphologies. An elegant alternative is the application of top-down lithographic procedures, which provides reproducible patterns of well-defined vertical nanostructures, such as nanopillars^16^, nanoneedles^17^ or nanocones^18^. However, the application of top-down strategies in biological studies is still constrained due to the requirement of complicated and limited-access technology such as focused ion beam (FIB) microscopy. In spite of these technological advances, a cost-effective method for substrate nanostructuration allowing for the quantification of how the external topography modulates the hierarchical organization and mechanical response of the cellular membrane is not yet developed. Here, we have combined a large-area lithographic method, such as Laser Interference Lithography (LIL), with the simplicity of soft-chemistry to engineer 1D SiO_2_ vertical nanostructures of defined sizes and shapes on high-performance coverslips suitable for advanced microscopies. These soft-gel nanoimprint lithography (soft-NIL) substrates, that we named soft-NIL templates, provides a large-area, customizable and benchtop equipment-based method to produce patterned nanostructures and manipulate the membrane morphology of living adherent cells. As a result of the combination of soft-NIL templates and advanced microscopies, here we show a quantification of the mechanical, chemical and dynamic state of cellular membranes in response to the morphology of the external environment. We provide insight on the topography-induced organization of proteins implicated in key cellular processes such as membrane trafficking, signaling and cell adhesion. This technological development of 1D nanostructured substrates provides a tool for several applications from fundamental mechanobiology to nanostructured bio-interfaces for tissue regeneration or bio-sensing devices.

## Results and discussion

### Soft-NIL templates deliver a customizable, large-area and cost-effective fabrication method to manipulate the shape of cellular membranes

To manipulate membrane shape in living cells we set up a large-area and cost-effective method to engineer 1D vertical nanostructures on high-performance coverslips made of borosilicate glass, which are the substrate of choice for super-resolution microscopy applications^19^. To this aim we customized Laser Interference Lithography (LIL)^20^ and NIL lithographic techniques, based on our previous work on epitaxial nanostructured α-quartz (SiO_2_) thin films^21,22^. Here, we adapted the quartz precursor solution^21^ to make dense and biocompatible nanostructured amorphous silica layers on glass substrates by removing the strontium devitrificant agent and the surfactants from the original solution. We combined the large-area nanostructuration capabilities of LIL, as compared to other elaborated top-down procedures, such as FIB or electron beam lithography (EBL), with bottom-up sol-gel chemistry to ultimately fabricate nanostructured SiO2 layers of controlled shape, diameter and periodicity. In a first top-down fabrication step, extensive Si(100) masters made of nanopillars arrays were obtained by using LIL and transferred by reactive ion etching at low pressure (see details in the methods section). A second step involved the preparation of high quality PolyDiMethylSiloxane (PDMS) molds from Si(100) masters (Figure S1). Next, we set up a dip-coating method to synthetize sol-gel silica films of controlled thicknesses on borosilicate glass coverslips in which we used the PDMS molds to produce perfectly imprinted silica nanopillars with controlled diameter and height on glass, as illustrated in Figure 1 and Figure S1. To achieve a perfect replication of silica pillars we optimized several parameters like temperature, humidity, thermal treatment or the hydrolysis degree of the silica sol, as detailed in the experimental section. Importantly, we found a strong dependence of the rate of pillar replication on the hydrolysis degree of the silica sol. Samples produced with a solution hydrolyzed more than 18 hours exhibited an incomplete transfer of the columnar pattern into the sol gel silica layer due to a rapid gelation process (as shown in Figure S2). As a result, we determined that the optimal interval for a perfect printing of the silica layer is 5 h to 16 h of hydrolysis process.

**Figure 1.**
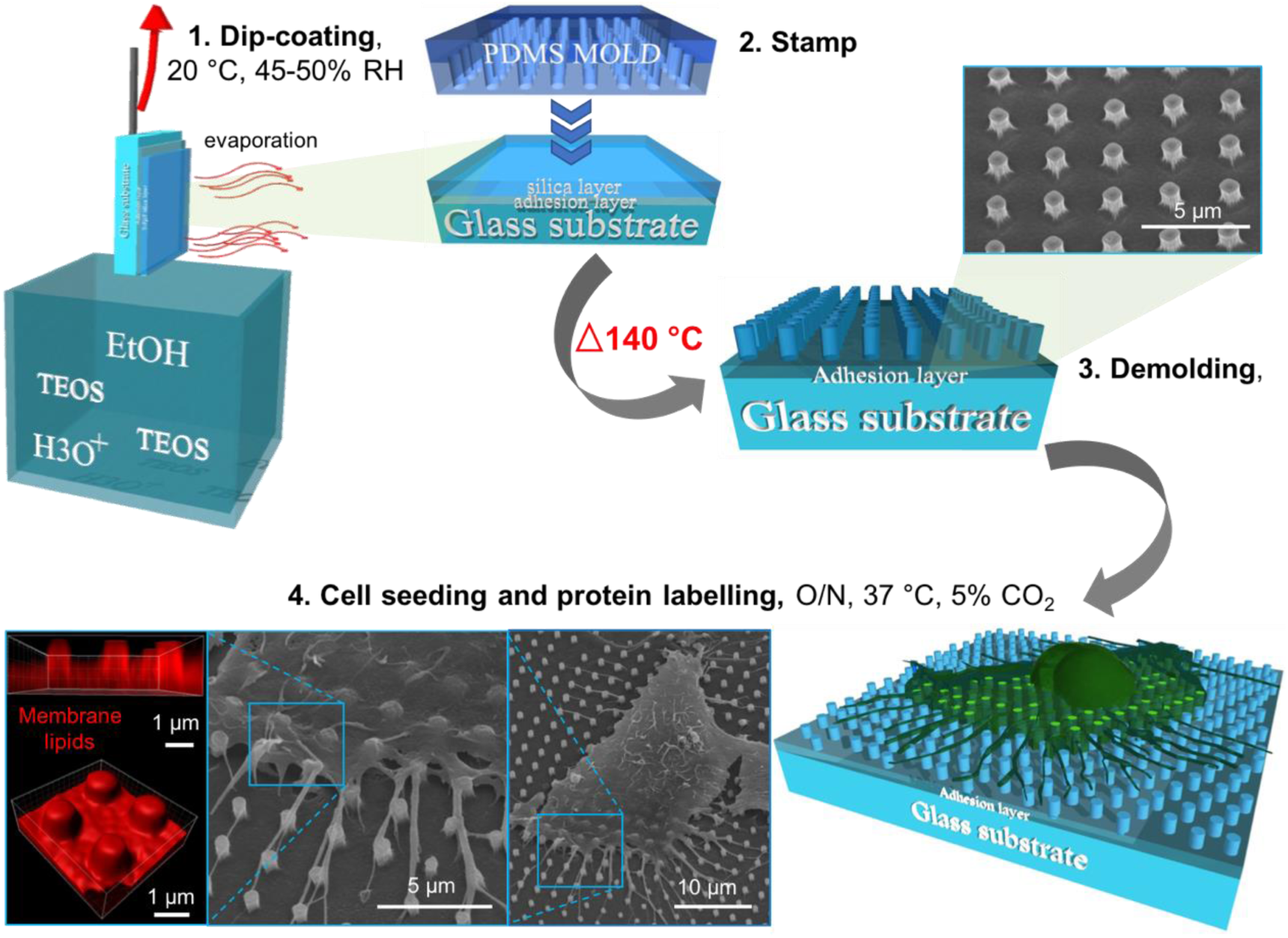
Schematic representation of the nanostructuration of SiO_2_ templates by soft-NIL to manipulate membrane shape in living cells. The main steps in the fabrication of 1D vertical SiO_2_ nanopillar arrays consisted in 1) dip-coating of a thin SiO_2_ sol-gel layer on high-performance coverslips, 2) stamping of the sol-gel layer with a PDMS mold, previously generated by LIL, 3) consolidation of the SiO_2_ sol-gel layer at high temperature (140°C) to create solid 1D vertical SiO_2_ nanostructures, as revealed by the SEM image. 4) Seeding of adherent cells on nanostructured SiO_2_ substrates allows to exert nanopillar-induced membrane deformations, as visualized by either SEM (*right and central panel*) or fluorescence microscopy of cell membrane lipids (*in red, left panel*).

Next, we evaluated whether our engineered nanostructured substrates, which we named “Soft-NIL templates”, were suitable to induce controlled plasma membrane deformations by seeding HeLa cells on vertical SiO_2_ nanocolumns of different sizes coated with poly-L-lysine. Parallel tests were also performed on C2C12 mouse myoblasts and the human breast cancer cell line MDA-MB-231. First, we confirmed the stability of silica nanopillars under cell culture conditions i.e. under typical cell culture medium at 37 ° C and 5% CO_2_. We found that 48 hours after cell seeding the pillar’s shape and dimension of soft-NIL templates remained intact all over the substrate surface. Bright-field images can be acquired on soft-NIL templates using either a 40X or 63X objective to visualize the arrays of pillars and seeded cells (Figure S3).

We confirmed that the cell plasma membrane was bent over the nanopillar structures as revealed from fluorescence microscopy images using a cell plasma membrane marker, the myristoylated and palmitoylated (MyrPalm) probe (Figure 1 and Figure S4), and by Scanning Electron Microscopy (SEM), as shown in Figure 1. Next, we evaluated the optical properties of soft-NIL templates for advanced microscopies. Imaging of supported lipid bilayers (SLBs) doped with different fluorescent lipid analogues showed that soft-NIL templates are suitable for the illumination of fluorophores using total internal reflection at highly inclined angles at the glass–medium interface (TIRF) and single point scanning microscopy (Airyscan)^19^ (Figure S5 and S6). We determined by Airyscan the fluorescence intensity of membranes on pillars (*I*_pillar_) respect to the fluorescence intensity away from pillars (*I*_basal_) using SLBs containing lipids conjugated to dyes emitting at three wavelengths: Oregon Green 488 DHPE (Ex/Em 501/526); TopFluor-TMR-PIP2 (Ex/Em 544/571 nm) and Atto647N DOPE (Ex/Em 643/662). In all cases, we obtained a ratio of I_pillar_/I_basal_ ∼ 1 and 1.3 in the case of Atto647N, suggesting that the pillar morphology itself should not induce, in principle, significant differences in the fluorescence signal of the conjugated dyes (Figure S6). Next, we compared the distribution of the cell membrane marker mCherry-MyrPalm with another cell marker, the Wheat Germ Agglutinin (WGA) conjugated to Alexa Fluor 488, with the fluorescent lipid analog rhodamine-DOPE using SLBs on soft-NIL templates displaying vertical nanopillars of different dimensions (Figure S6). We systematically obtained a ratio of I_pillar_/I_bottom_ ∼ 1 for the three membrane markers and different nanopillar dimensions, suggesting that, under these experimental conditions, the distribution of membrane markers at the cell surface is not substantially affected. Altogether these results indicate that soft-NIL templates constitute a tool to manipulate membrane shape on adherent living cells.

As a result, we designed different silica patterns to induce controlled plasma membrane deformation in live cells. Silica nanostructured substrates were composed of vertical nanopillars of well-defined diameter, height, period and shape, i.e. a diameter of 400 and 600 nm for the circular pillars and a diagonal of 800 nm for the square nanopillars (as shown in Figure 2a-c and detailed in table 1 of the supporting information). Because the production of patterns by LIL is determined by the diffraction limit of the technique, i.e. half of the laser beam wavelength, we set up an original method to reduce the dimensions of silica nanopillars in a controlled manner. This approach consists in producing PDMS molds from each sol-gel replica successively as illustrated in Figure 2d and e, thus reducing the silica nanopillar’s dimension of 40% at each NIL process, which corresponds to a reduction of around 200 nm in height from the original silicon master after consolidation at 430 °C during 10 min.

**Figure 2.**
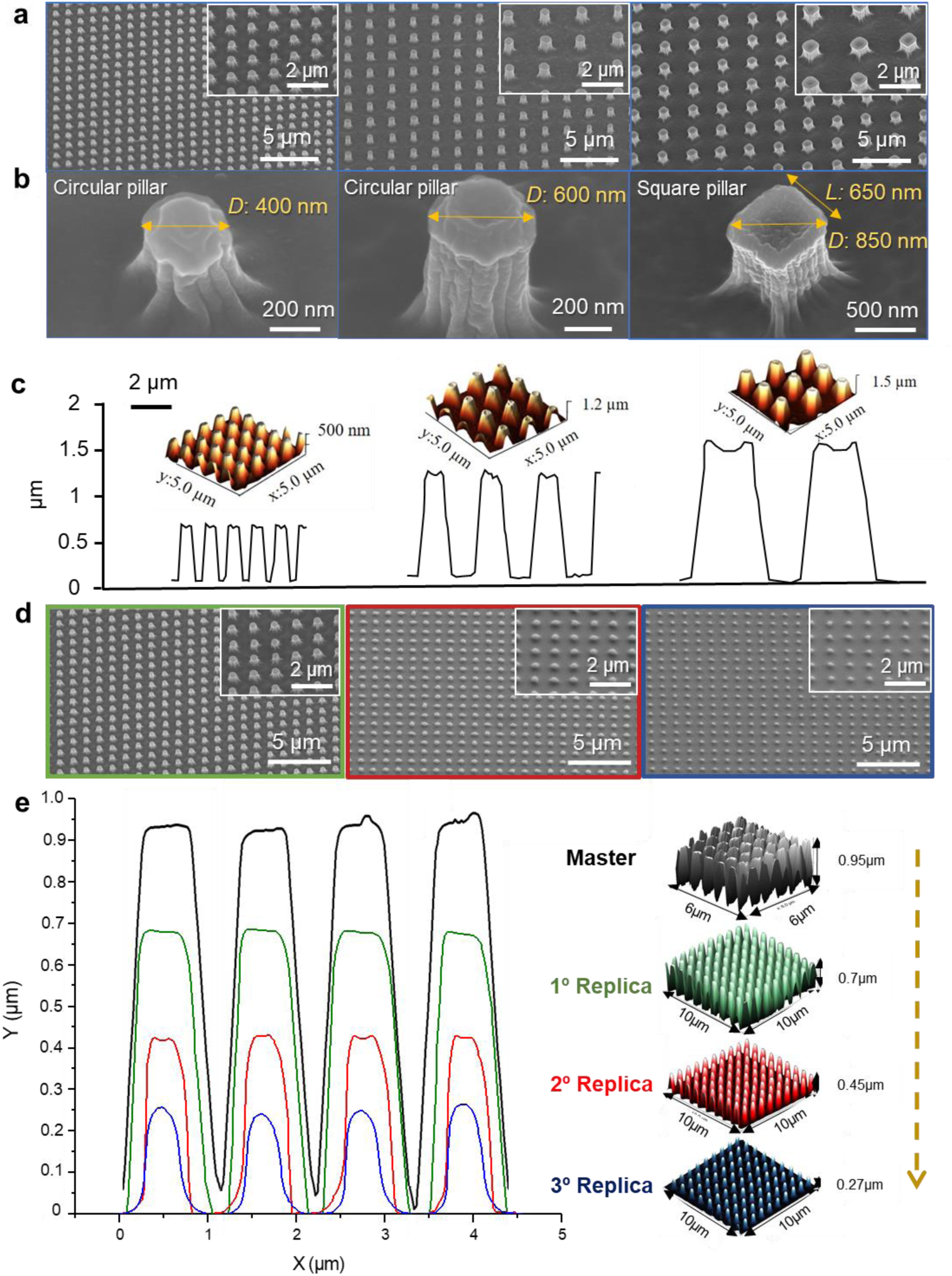
Control of vertical SiO_2_ nanocolumn dimensions and geometry by a multiple replication process of the original silicon master. **a**, FEG-SEM images of printed silica nanopillars with controlled diameter i.e. 400 and 600 nm of diameter for the circular pillars and 800 nm of diagonal for squared nano pillars. **b**, FEG-SEM images at higher magnification of pillars showing the different dimensions. **c**, 3D AFM images showing silica nanostructured films prepared by NIL lithography in a. Below you can distinguish the profile analysis of the AFM image in a, revealing a perfect transfer of the different motifs. **d**, FEG-SEM images of printed silica nanopillars with reduced dimension by multiple soft-NIL replicates. **e**, Notice that after applying this method the silica nanopillar reduces their heights by 40% in each NIL process approximately, which corresponds to a reduction in height of around 200 nm from the original silicon master (*in black*), as shown by the corresponding AFM topography profiles.

### Soft-NIL templates remodel the chemical and physical state of the plasma membrane

Next, we evaluated the capabilities of soft-NIL templates to modulate the physical and chemical state of the plasma membrane by determining the plasma membrane tension resulting from seeding HeLa cells on soft-NIL templates of dimensions (*D*) *D* ∼ 400 nm, *D* ∼ 600 nm and *D* ∼ 850 nm. To this end, we applied the fluorescent membrane tension sensor FliptR, which provides a direct estimation of the membrane tension and/or lipid packing of living cells by using fluorescence lifetime imaging microscopy (FLIM), as recently reported^23^. As shown in Figure 3a-c, we found that the mean FliptR lifetime was systematically higher at the cell membrane localized at the base of pillars, here denoted as basal membrane, compared to the membrane bent over the pillars, here denoted as on pillars, irrespective of the soft-NIL template dimension. Whereas a global decrease of the mean FliptR lifetime was observed between pillars of *D* ∼ 400 nm and *D* ∼ 600 nm with τ = 4.80 ± 0.11, τ = 4.20 ± 0.34 and τ = 4.61 ± 0.13, τ = 4.05 ± 0.38 (mean ± s.d.) for the basal membrane and the membrane on pillars, respectively, we did not observe a significant change on soft-NIL templates of *D* ∼ 600 nm and *D* ∼ 850 nm (Figure 3c). Thus, indicating that the morphology and periodicity imposed by soft-NIL templates might affect the tension and/or lipid packing at the pillar-cell membrane interface. Next, we investigated whether the dimension and periodicity soft-NIL templates could also trigger local changes in the distribution of lipids on cellular membranes by assessing the localization of the main plasma membrane phosphoinositides (PIs) – phosphatidylinositol 4,5-bisphosphate [PI(4,5)P2] and phosphatidylinositol 3,4,5-trisphosphate [PI(3,4,5)P3] - relative to the plasma membrane marker MyrPalm. Indeed, the localization of both PI(4,5)P2 and PI(3,4,5)P3 at the plasma membrane is required for the precise recruitment of many signaling, cytoskeletal and trafficking proteins, such as small GTPases, via electrostatic interactions^24^. We detected the endogenous localization of both PIs species on HeLa cells seeded on soft-NIL templates of *D* ∼ 400 nm, *D* ∼ 600 nm, *D* ∼ 850 nm by using fluorescent recombinant PI-binding probes^25^. Our results show that soft-NIL templates of *D* ∼ 400 nm engage a moderate but local accumulation of both PI(4,5)P2 and PI(3,4,5)P3 at the plasma membrane-pillar interface with an average 1.3 ± 0.3 and 1.3 ± 0.4 (mean ± s.d.) fold increase, respectively, as compared to the MyrPalm probe (Figure 3d). Collectively, our results suggest that the membrane morphology denoted by soft-NIL templates might affect the distribution of relevant lipids such as the PIs at the plasma membrane (Figure 3d), although experiments would need to be conducted to determine if the accumulation of PI’s on vertical nanopillars is able to trigger the recruitment of PI-interacting proteins and/or the activation of PI-mediated signaling pathways.

**Figure 3.**
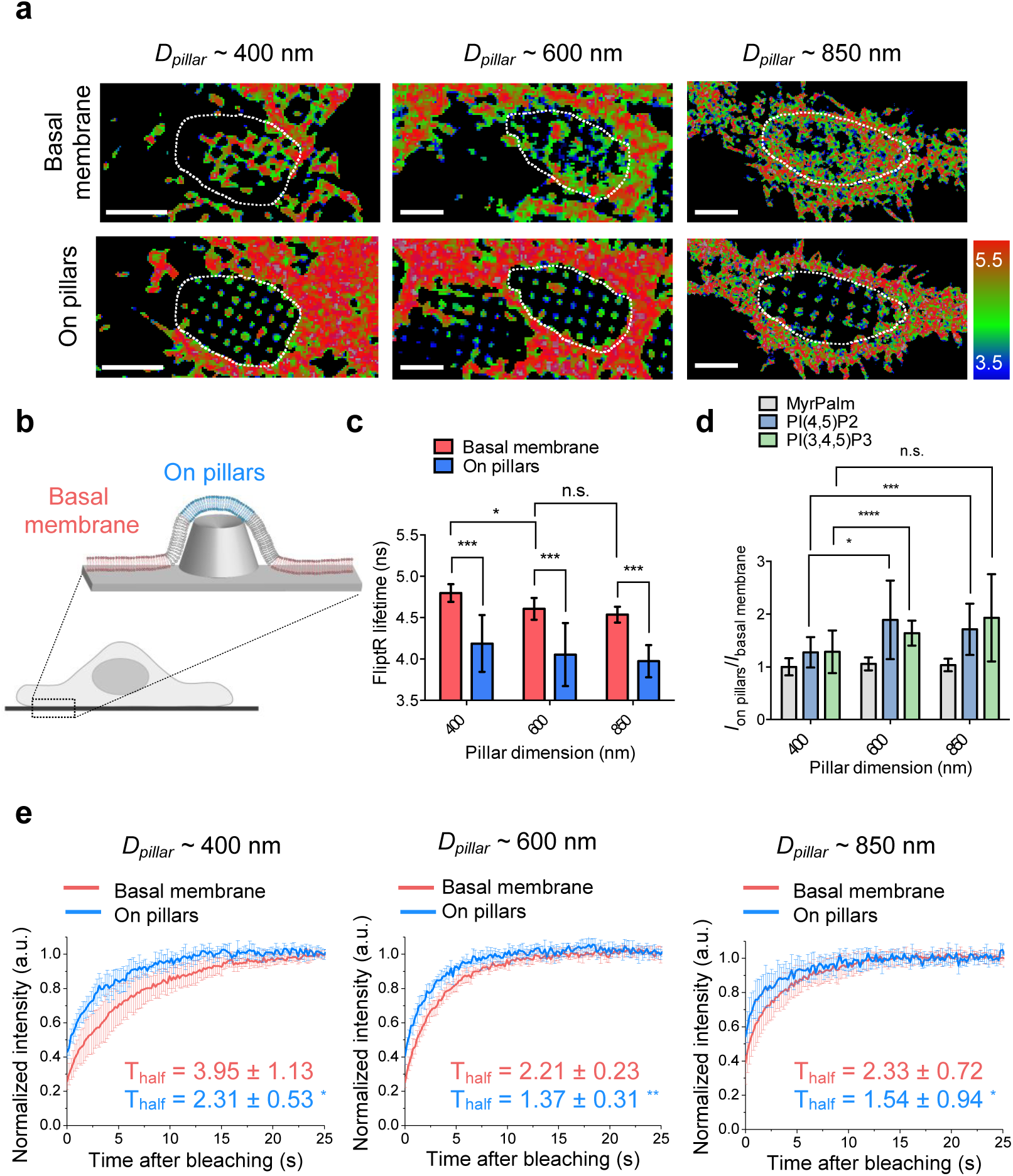
1D vertical nanopillars modify the membrane tension and the lateral organization of the plasma membrane. **a**, Representative FLIM images showing the FliptR lifetime (ns) of two confocal planes (at the basal membrane, i.e. at the bottom of pillars, *top panel*, and on pillars, i.e. at the top of pillars, *bottom panel*) of the plasma membrane of HeLa cells seeded on vertical nanopillars of *D* ∼ 400 nm, *D* ∼ 600 nm, and *D* ∼ 850 nm and stained with the membrane tension probe FliptR. The regions of interest that were used for the data analysis are highlighted in each image. Notice that confocal planes acquired at the top of pillars also display the FliptR staining of cell membrane regions away from pillars, e.g. between adjacent cells. The color bar code corresponds to lifetime in ns. Scale bar, 2 μm. **b**, Schematics of the membrane regions of interest: the plasma membrane in contact with the flat surface of soft-NIL templates, denoted as “basal membrane”, and the plasma membrane bent over vertical nanopillars on soft-NIL templates, denoted as “on pillars”. **c**, FliptR lifetime (ns) acquired at basal membrane (*red*) and at the membrane-pillar interface (*blue*) on HeLa cells seeded on soft-NIL templates of *D* ∼ 400 nm, *D* ∼ 600 nm, and *D* ∼ 850 nm. \**P*<0.03, \*\*\*\**P*<0.0001 (two-way ANOVA). Error bars represent s.d.; *n*=13, *n*=144 for *D* ∼ 400 nm; *n*=14, *n*=221 for *D* ∼ 600 nm; *n*=16, *n*=96 for *D* ∼ 800 nm, for the basal membrane and on pillars, respectively. **d**, Ratio of the intensity of different plasma membrane lipid probes measured on pillars relative to the basal membrane as a function of soft-NIL templates dimension for MyrPalm (*gray*), PI(4,5)P2 (*blue*), and PI(3,4,5)P3 (*green*). Dotted gray line indicates 1. \*\*\*\**P*<0.0001 (two-way ANOVA). Error bars represent s.d.; *n*=691, *n*=295, *n*=152 for MyrPalm; *n*=218, *n*=345, *n*=161 for PI(4,5)P2; *n*=82, *n*=70, *n*=80 for PI(3,4,5)P3 for *D* ∼ 400 nm, *D* ∼ 600 nm, and *D* ∼ 800 nm, respectively. **e**, FRAP curves showing the membrane diffusion, approximated by the T_half_ value at the basal membrane. Each curve represents the average fluorescence over time. \**P*<0.03, \*\**P*=0.003 (Paired t-test). Error bars represent s.d.; n > 5.

Because the local organization of membranes is often associated to changes in the dynamic behavior of its building components^26^, we set out to investigate the lateral diffusion of the plasma membrane by performing Fluorescence Recovery After Photobleaching (FRAP) measurements on HeLa cells expressing the MyrPalm probe and seeded on soft-NIL templates of different dimensions (see Figure S7 for a representative FRAP acquisition). As shown in Figure 3e, our results revealed a systematic increase in the FRAP times at the membrane-pillar interface (i.e. on pillars) as compared to the basal membrane (i.e. at the base of vertical nanopillars) irrespective of the nanopillar dimensions, in agreement with the trend observed when using the lipid tension reporter FliptR (Figure 3c). Indeed, the obtained half-time of recovery (T_half_) values were globally higher on cells seeded on soft-NIL templates of *D* ∼ 400 nm as compared to *D* ≥ 600 nm, suggesting that the nanopillar dimension and periodicity might affect the lateral organization of cellular membranes. Our results show that membrane tension and FRAP times display a sustained behavior that is distinct between the longitudinal axis of nanopillars, as denoted by the decrease in the mean lifetime of the tension probe FliptR and a faster FRAP on top of the pillars as compared to the basal membrane (Figure 3).

### Mapping of the steady-state organization of cell surface receptors using soft-NIL templates

Next, we assessed weather the distinct mechanical and chemical response of cellular membranes on soft-NIL templates could be transduced in changes in the steady-state organization of transmembrane receptors at the cell surface. To quantitatively determine the tendency of cellular receptors to locally accumulate on membrane morphologies denoted by vertical nanostructures we estimated the receptor’s local enrichment, *E*, which stands for the ratio of the receptor intensity on pillars *versus* the basal membrane and normalized by the MyrPalm probe, as described in the methods section. We considered that an *E* larger than 1 would reflect a preferential localization at the plasma membrane-nanopillar’s interface with respect to the basal membrane away from the nanopillars. By using Airyscan microscopy, we monitored the localization of different receptors on HeLa cells seeded on soft-NIL templates of *D* ∼ 400 nm and *D* ∼ 600 nm. To this aim, we used HeLa cells expressing either the transferrin receptor (TfR), which is a nonsignaling receptor constitutively internalized through clathrin-coated pits, or the signaling epidermal growth factor receptor (EGFR), which is a tyrosine kinase receptor^27,28^. In addition, we monitored by immunostaining the endogenous steady-state localization of the β1-integrin receptor, which is the β-subunit of the fibronectin and collagen-binding integrin heterodimeric receptor mediating the adhesion of cells with the extracellular matrix (ECM)^29^. As displayed in Figure 4a-c, we observed an accumulation of the TfR, EGFR and β1-integrin on vertical nanostructures with a moderate but significant difference between the two pillar dimensions in the case of TfR, whereas this was not the case for the EGFR and β1-integrin. The different enrichment of the TfR on vertical nanopillars of *D* ∼ 400 nm and *D* ∼ 600 nm might indicate that its accumulation might respond, either directly or indirectly, to the morphology of membranes and possibly, enhanced by the slower FRAP recovery observed in small pillar sizes as compared to the larger sizes (*D* ≥ 600 nm) (Figure 3e). We observed that the recruitment of endocytic proteins holding a curvature sensing domain, such as the F-BAR protein FCHo2 was also enhanced on pillars of *D* ∼ 400 nm respect to larger sizes, *D* ≥ 600 nm (Figure S8). Both AP-2 and FCHo2 are endocytic proteins implicated in the initiation of the clathrin pathway^30^ and appear concomitantly enriched on vertical nanopillars, particularly at *D* ∼ 400 nm (Figure S9). Thus, an increased accumulation of endocytic proteins at the membrane-pillar interface might explain the enrichment of clathrin-regulated surface receptors on vertical nanostructures. Our observations using the sensor probe FliptR showed an inverse relationship between pillar dimension and plasma membrane tension (Figure 3a-c). Thus, it might be also possible that by stabilizing a membrane shape that is compatible with curvature sensing proteins, vertical nanostructures of *D* ∼ 400 could by-pass the inhibitory effect that increased membrane tension has in clathrin-mediated endocytosis^31^.

**Figure 4.**
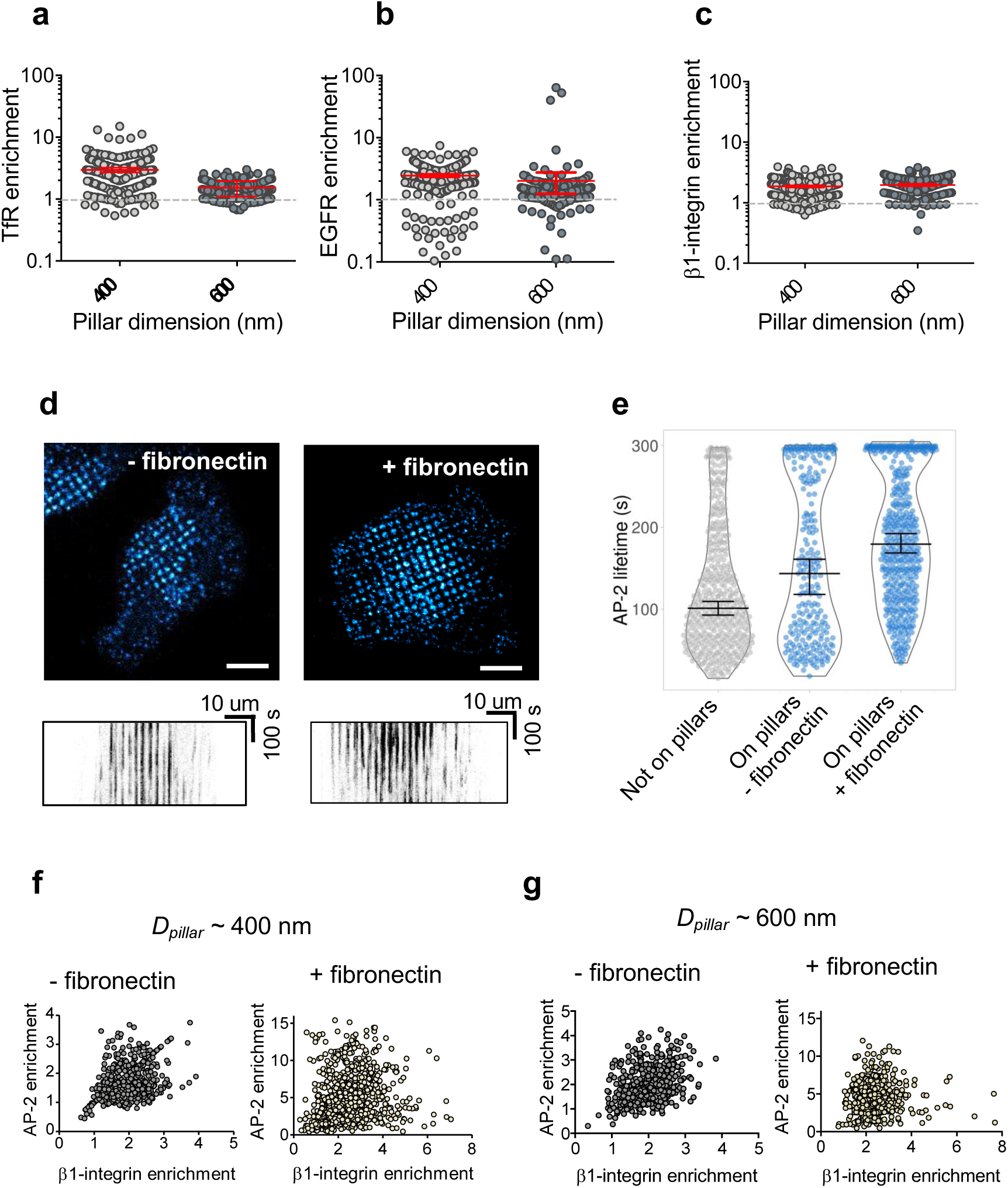
Steady-state organization of cell surface receptors on 1D SiO_2_ nanopillar’s arrays. **a-c**, Enrichment, *E*, of TfR, EGFR and β1-integrin on HeLa cells seeded on nanopillars of *D* ∼ 400 nm (*gray*) and *D* ∼ 600 nm (*slategray*). Data points per condition are: *n*=222, *n*=214 for TfR; *n*=332, *n*=231 for EGFR; *n*=332, *n*=234 for β1-integrin, for the nanopillar’s of *D* ∼ 400 nm and *D* ∼ 600 nm, respectively. Dotted grey line indicates an *E* ∼ 1. **d**, The dynamics of AP-2 was obtained by spinning disk microscopy every 2s for 5 min on genome edited AP-2 s2-EGFP SUM159 cells seeded on soft-NIL templates of *D* ∼ 600 nm either coated with poly-L-lysin (-fibronectin) or fibronectin (+ fibronectin). Still images represent maximum intensity projections of z-stacks at t = 0s. Representative kymographs are shown below each corresponding condition. **e**, Lifetime (in s) of AP-2 from s2-EGFP SUM159 cells seeded on conventional SiO_2_ coverslips coated with poly-L-lysin (not on pillars, *in gray*) or on soft-NIL templates of *D* ∼ 600 nm (on pillars, *cadetblue*) either coated with poly-L-lysin (-fibronectin) or fibronectin (+ fibronectin), as observed in the kymographs in **d**. The number of data points per condition are *n*=440 (not on pillars), *n*=255 (-fibronectin) and *n*=461 (+ fibronectin). **f-g**, Scatter plots showing enrichment of β1-integrin and AP-2 per pillar on SUM159 cells seeded on soft-NIL templates of *D* ∼ 400 nm and *D* ∼ 600 nm coated (+ fibronectin, *light yellow*) or not (-fibronectin, *gray*) with fibronectin. The number of data points per condition are: *n*=625 and *n*=844 for *D* ∼ 400 nm; and *n*=426 and *n*=500 for *D* ∼ 600 nm for the non-coated and fibronectin coated substrates, respectively. The obtained correlation coefficient for each condition is: r = 0.41, r = 0.18, r = 0.37 and r = 0.04, respectively.

It is reported that signaling receptors, such as the EGFR, cluster as a consequence of ligand binding. Although the relative contribution of receptor clustering in the initiation of signaling and endocytosis is not well understood, we wonder if the accumulation of EGFR that we observed on soft-NIL templates in the absence of EGF might affect the receptor’s activity. To this end, we estimated by immunostaining the apparent phosphorylation of the EGFR at Tyr1045, which creates a major docking site for the c-Cbl adaptor protein leading to receptor ubiquitination after EFGR activation^32^. We found a mean *E* ≥ 3 of phospho-EGFR (Tyr1045) antibody on HeLa cells seeded on soft-NIL templates of dimensions *D* ∼ 400 nm and *D* ∼ 600 nm (Figure S10).

The accumulation of the endocytic protein AP-2, which has been associated to signaling and adhesion receptors^33,34^, in vertical nanostructures of D ≥ 400 nm (Figure S9) let us investigate if other parameters than membrane curvature might contribute to its surface organization. A parameter that could be affected by the pillar morphology is membrane dynamics, as observed in Figure 3. Previous works reported that on HeLa cells two populations of AP-2/clathrin structures co-exist at the ventral surface of seeded cells: small and dynamic “coated pits” and larger and long-lived “coated plaques”^35^. While the dynamics of pits and plaques is often discriminated on cells seeded on flat surfaces due to their characteristic brightness^33^, the limited resolution to distinguish these structures at different z-positions along the nanopillar let us consider another cell type to investigate the AP-2 dynamics. The absence of long-lived AP- 2/clathrin structures reported on gene-edited SUM159-AP-2 cells expressing σ2-EGFP^36^ provides a convenient model to examine the effect of vertical nanostructures on AP-2 dynamics, either by affecting the lifetime or the initiation rate of, in principle, coated pits. We could confirm that gene-edited SUM159-AP-2 cells expressing σ2-EGFP also presented a ∼2- fold enrichment of AP-2 on membrane shapes with radii > 200 nm, although to a lesser extend to that observed in HeLa cells (Figure S9). As shown in Figure 4d-e, we found an increase in the average lifetime of AP-2 structures on soft-NIL templates, as compared to conventional SiO_2_ substrates. Integrins are one of the major families of cell adhesion receptors and coating of soft-NIL templates with fibronectin and collagen, which are integrin ligands, also increased AP-2 lifetime (Figure 4d-e and Figure S11). The engagement and accumulation of integrins upon tight binding with the extracellular matrix has been described to hinder the lateral diffusion of membrane components^37,38^, which strongly depends on the molecular crowding but also on other parameters such as the local composition and the actin cytoskeleton^26^. Therefore, explaining the increase in the local enrichment of AP-2 structures on fibronectin-coated soft-NIL templates, as compared to non-coated substrates observed in Figure 4f-g and Figure S12. We found that in the absence of fibronectin the enrichment of AP-2 and β1-integrin display a moderate correlation, with a correlation coefficient of r = 0.41 and r = 0.37 on nanopillars of D ∼ 400 nm and D ∼ 600 nm, respectively. Because integrin cytoplasmic tails were shown to interact with clathrin adaptors such as Dab2 and ARH that bind to β-tail motifs and the μ2- subunit of the AP-2 that binds to a subset of α-subunit tails^39,40^ during clathrin-mediated endocytosis, this interaction might explain the correlation that we observed between AP-2 and β1-integrin on vertical nanopillars (Figure 4f-g). However, addition of fibronectin reduced the correlation between AP-2 and β1-integrin on vertical nanopillars, with a r = 0.18 and r = 0.04 on nanopillar’s of D ∼ 400 nm and D ∼ 600 nm, respectively. Thus, indicating that β1-integrin engagement might not participate in the accumulation of AP-2 and possibly, its dynamics, on vertical nanostructures and that other integrins could be involved in this process. This observation is in agreement with previous works reporting that the formation of long-lived AP- 2/clathrin structures is convoyed with the enrichment of signaling receptors and integrins^33,34^. Establishing how membrane shape affects the lateral organization and activity of transmembrane receptors at the cell surface is crucial to understand many cellular functions^41^. Although seminal works have successfully showed the effect of curvature on the receptor sorting *in vitro*^42,43^, these studies often require complicated reconstitution procedures. Our results show that membrane shape has an active role assisting the steady-state organization and dynamics of protein complexes at the cell surface, a process that can be investigated using soft- NIL templates.

### Soft-NIL templates assist the formation of protein-specific territories at the plasma membrane

Next, we set out to investigate if the membrane properties along vertical nanopillars might also assist the steady-state organization of protein complexes at the cell surface. To this aim, we monitored by Airyscan imaging the distribution of the surface receptors TfR, EGFR and β1- integrin as well as the endocytic proteins involved in their cellular uptake and the actin cytoskeleton on HeLa cells seeded on soft-NIL templates of either *D* ∼ 400 nm or *D* ∼ 600 nm (Figure 5). As shown by the representative 3D visualization in Figure 5 a-b, the membrane lipid PI(4,5)P2 is localized all along the membrane wrapping the nanopillars and thus, renders a visualization of the pillar contour. The distribution of endogenous clathrin and AP-2 obtained by immunostaining displays a discrete signal along the z-axis of vertical nanopillars. This is not the case of the F-actin staining, which is often localized at the base of nanopillars. We estimated the maximal height of the AP-2, clathrin and actin signal relative to the PI(4,5)P2 signal, which defined the position of the columns and their heights, as detailed in the methods section. The representative cross-section of the resulting maximal heights, as shown in Figure 5b, also suggests that AP-2, clathrin and actin display a specific surface distribution along the z-axis of soft-NIL templates, while F-actin is frequently excluded from the top of pillars (Figure 5a-b). The frequency distribution of the maximal height obtained for actin, AP-2 and clathrin on soft-NIL templates of *D* ∼ 400 nm and of *D* ∼ 600 nm is shown in Figure 5c. We found an average maximal height of 0.71 ± 0.26, 0.87 ± 0.16, and 0.86 ± 0.17 (in a.u., mean ± s.d.) for actin, AP-2 and clathrin, respectively, on soft-NIL templates of *D* ∼ 400 nm. Their specific distribution was accentuated on soft-NIL templates of *D* ∼ 600 nm, which presented an average maximal height of 0.58 ± 0.29, 0.94 ± 0.13, 0.96 ± 0.11 (in a.u., mean ± s.d.) for actin, AP-2 and clathrin, respectively. Collectively, indicating that soft-NIL templates reveal a specific distribution of membrane-associated proteins along the z-axis of vertical nanostructures. Because we found that soft-NIL templates facilitate the local enrichment of the surface receptors TfR, EGFR and β1-integrin at the plasma membrane-pillar interface, as shown in Figure 4a-c, we estimated their surface distribution relative to the position of the endocytic proteins implicated in their uptake, i.e. AP-2/clathrin structures (Figure 5d). We found that the TfR is preferentially distributed at the upper region of vertical nanopillars with an average maximal height of 0.96 ± 0.08 (in a.u., mean ± s.d.). This was also the case of the EGFR and β1-integrin signals, but with a distribution that was moderately skewed towards the bottom of pillars as compared to the TfR, with a maximal height of 0.89 ± 0.17 and 0.84 ± 0.16 (in a.u., mean ± s.d.), respectively. Thus, in spite of the fact that TfR, EGFR and β1-integrin are all internalized through the clathrin pathway, our results suggest that each surface receptor might display a preferential distribution on membrane morphologies denoted by vertical structures of *D* ∼ 400 nm.

**Figure 5.**
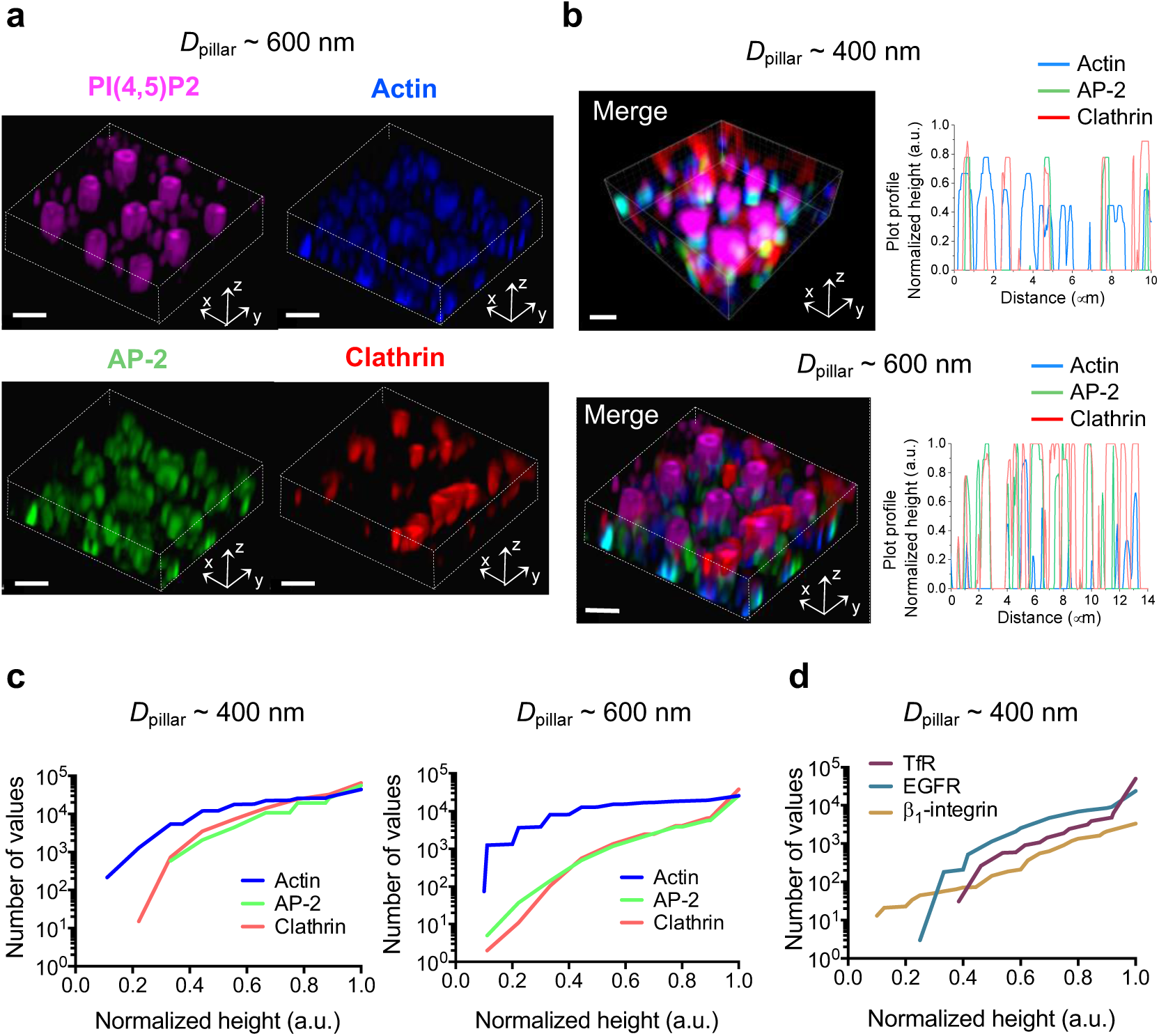
Soft-NIL templates assist the surface localization of cellular proteins along the z-axis of vertical nanostructures. **a**, Representative 3D reconstruction of the plasma membrane of HeLa cells seeded on nanopillars of *D* ∼ 600 nm showing the distribution of the lipid PI(4,5)P2 (*magenta)*,which highlights the contour of the membrane, actin (*blue*), AP-2 (*green*) and clathrin (*red*). Scale bar, 1 μm. **b**, Representative 3D image of the plasma membrane of HeLa cells seeded on nanopillars of *D* ∼ 400 nm and *D* ∼ 600 nm showing the co-localization (merge image) between PI(4,5)P2 (*magenta*), actin (*blue*), AP-2 (*green*) and clathrin (*red*) along the z-axis of vertical nanopillars. Scale bar, 1 μm. Representative profile analysis of the maximal heights of actin (*blue*), AP-2 (*green*), clathrin (*red*) normalized by the pillar height denoted by the PI(4,5)P2 signal for each of the two pillar dimension, *D* ∼ 400 nm and *D* ∼ 600 nm. **c**, Cumulative frequency distribution of the maximal heights obtained for actin (*blue*), AP-2 (*green*) and clathrin (*red*) and normalized by the pillar height on HeLa cells seeded on soft-NIL templates of *D* ∼ 400 nm and *D* ∼ 600 nm. *n*=44182, *n*=55706, *n*=65535 for *D* ∼ 400 nm; *n*=25388, *n*=25740, *n*=38130 for *D* ∼ 600 nm and for the actin, AP-2 and clathrin signals, respectively. **d**, Cumulative frequency distribution of maximal heights obtained for TfR (*purple*), EGFR (*cyan*) and β1-integrin (*orange*) and normalized by the pillar height on HeLa cells seeded on soft-NIL templates of *D* ∼ 400 nm. *n*=50286, *n*=23774, *n*=3362 for the TfR, EGFR and β1- integrin signals, respectively.

### Soft-NIL templates support a hierarchical organization of the cell membrane

Next, we investigated whether the specific distribution of membrane-associated proteins, as observed in Figure 5, might display a discrete or long-range organization on soft-NIL templates. To this aim, we determined the 2D autocorrelation of the actin, AP-2 and clathrin staining on HeLa cells seeded on two soft-NIL template morphologies, circular and square shape, as shown by the high-magnification of SEM images in Figure 6. The structural assembly of the three proteins into long-range 2D periodic lattices revealed a discrete self-organization around the nanopillars irrespective of the pillar’s shape, an organization that we did not observe on SiO_2_ flat substrates (Figure S13). In the case of AP-2 and clathrin we observed an isotropic organization around circular pillars and a remarkable anisotropic arrangement on square nanopillars (Figure 6c and f). The average distances obtained between two maxima of the repetitive elements in the normalized surface autocorrelation function corresponding to fluorescence images, as displayed in Figure 6b and e, showed a periodic distribution of 1086 ± 70 nm, 1047 ± 50 nm, and 1041 ± 60 nm for cells seeded on circular pillars, and 2409 ± 4 nm, 2374 ± 10 nm and 2405 ± 20 nm (mean ± s.d., *N ≥ 5*) for cells seeded on square pillars, for actin, AP-2 and clathrin, respectively, along the spatial directions [010] and [100] of vertical nanostructures (Figure 6c and f). The long-range patterning obtained for each of the actin, AP- 2 and clathrin signals agreed with the periodicity of the corresponding soft-NIL templates (Table 1 in supporting information). Thus, indicating that the structural arrangement of proteins at the cell surface is imposed by the morphology and periodicity of soft-NIL templates. However, each protein complex that we tested in this study displayed a particular hierarchical patterning along the three spatial directions of vertical nanostructures, i.e. [100], [010] and [001] directions, (Figure 4 and Figure 5). This singular structural arrangement of proteins that we observed on soft-NIL templates might reflect the different modes of association with and within cellular membranes, for instance as a result of the ability to form semi-flexible polymers, such as actin^44^, or to oligomerize into supra-molecular complexes, such as clathrin lattices^45^.

**Figure 6.**
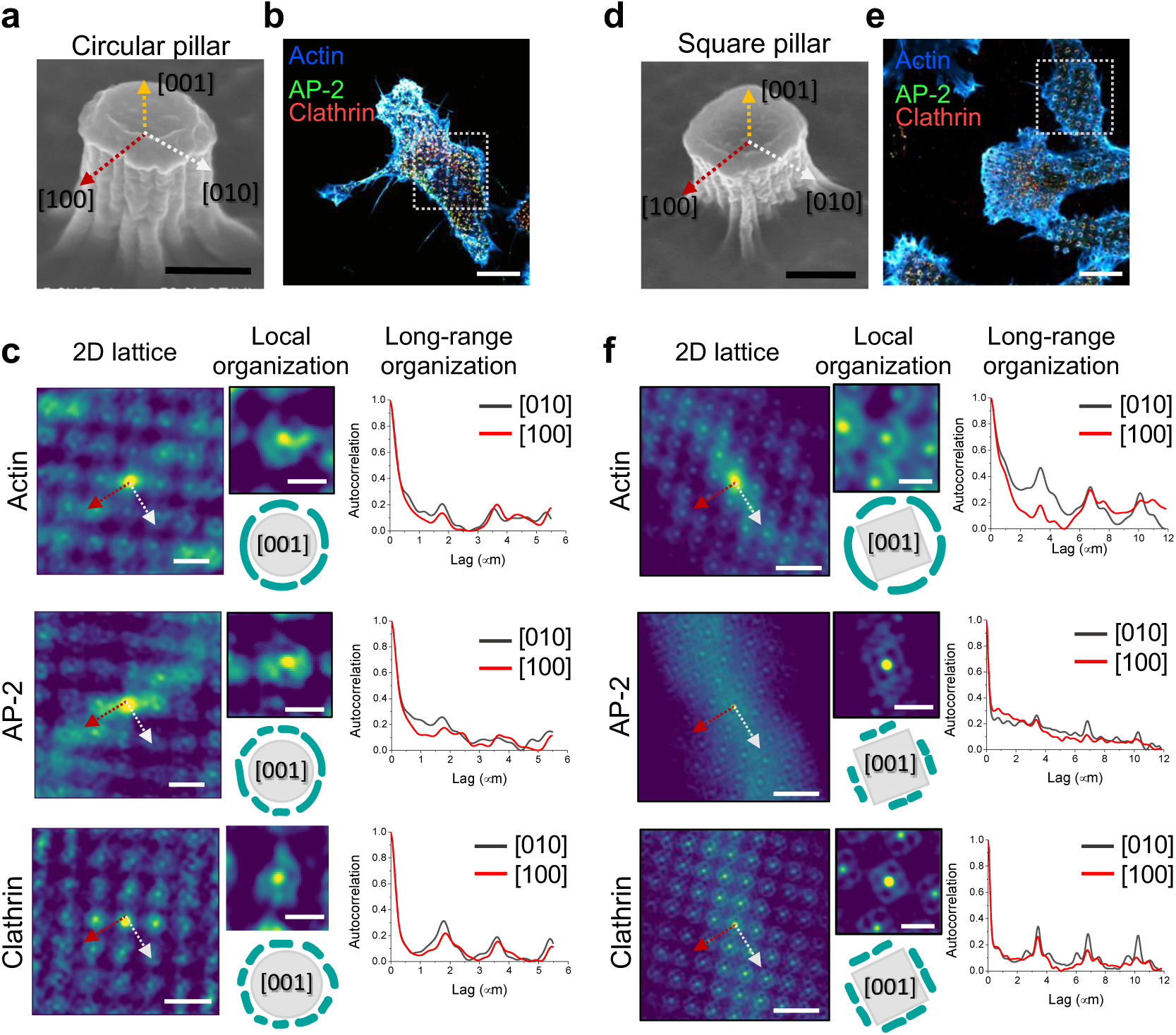
Vertical nanostructures assist the local and long-range self-organization of proteins at the cell membrane. **a**, High-magnification SEM image of a circular nanopillars. Scale bar, 400 nm. **b**, Maximum intensity projection of an Airyscan image of HeLa cells seeded on circular nanopillars and immunostained for actin (*blue*), AP-2 (*green*), and clathrin (*red*). Scale bar, 10 μm. **c**, 2D autocorrelation function of actin, AP-2, and clathrin signals corresponding to the white dashed box in **b**. 4X magnifications of the corresponding 2D-lattice to reveal the local structural arrangement of each protein signal. Scale bar, 2 μm. Schematic representation of the self-organization of actin, AP-2, and clathrin signals (*cyan*) around vertical nanopillars (*gray*), as observed in the corresponding 2D lattice. Normalized surface autocorrelation function along the crystallographic directions [010] and [100] of vertical nanostructures, as indicated by the red and gray arrows in the corresponding image. **d-f**, same as a-c, but on rectangular nanopillars.

### Conclusions

In conclusion, we showed the applicability of soft-NIL for the simple and cost-effective nanostructuration of high-performance glass compatible with advanced microscopies to investigate the transduction of topological features at the plasma membrane of live cells. Thus, providing an alternative to the limited access of FIB technology in cell biology and biophysical studies. By using soft-NIL templates we analyzed how vertical SiO_2_ nanostructures affect the mechanics and dynamic organization of cellular membranes. We showed that on patterned surfaces the local and long range organization of cell surface receptors, such as the EGFR, TfR and β1-integrin, as well as the proteins implicated their uptake is modulated by the shape of cellular membranes. Because the formation of specialized domains on membranes is a relevant feature to support signaling, membrane trafficking, cell adhesion, and host-pathogen interactions^24,46^, soft-NIL templates constitute a tool to investigate these processes in a controlled manner *in cellulo*. The biocompatibility of soft-NIL templates also opens the door to biotechnical applications including the development of biosensors.

## Materials and Methods

### Synthesis of silica sol-gel thin films

#### Solution preparation

All the chemicals were from Sigma-Aldrich and without any further purification. Silica precursor solution were prepared by adding 4.22 g tetraethyl orthosilicate (TEOS) into 23.26 g absolute ethanol, then 1.5 g HCl (37%), and stirring the solution for 18 h. The final molar composition was TEOS:HCl:EtOH=1:0.7:25.

#### Gel films by dip-coating

layer gel films on high-performance coverslips made of borosilicate glass of the first hydrolytic class with precision thickness No. 1.5 (0.170 ± 0.005 mm) at room temperature were prepared with a ND-DC300 dip-coater (Nadetech Innovations) equipped with an EBC10 Miniclima Device to control the surrounding temperature and relative humidity. During the dip-coating, we fixed the ambient temperature and relative humidity as 20ºC and 45-50% and the thickness of film was controlled by the withdrawal rate. In this study, all the films were made at withdrawal rate of 300 mm/min i.e. 200 nm thick to ensure the nanoimprint process. After dip-coating, as-prepared gel films were consolidated with a thermal treatment of 5 min at 430 ºC under air atmosphere.

### Silica thin film nanostructuration

#### Molds preparation

Si masters were elaborated using LIL Lithography as previously reported^21,22^. Briefly, this procedure allows to quickly obtain periodic patterns on silicon substrates of controlled shape (square and circular, as previously described^47^, diameter and periodicity over a large surface (∼cm^2^) without the need of a lithographic mask^20^.

PDMS (polydimethylsiloxane) reactants (90 w% RTV141A; 10 w% RTV141B from BLUESIL) were transferred onto the master and dried at 70 °C for 1 h before unmolding. Then, a first silica layer seed was deposited at a constant relative humidity of 45-50% at 20ºC with controlled withdrawal speeds of 300 mm min^−1^ in order to adjust the final thickness to 200 nm, and was consolidated at 430ºC for 5 min. Importantly, this film has an important functionality as an adhesion layer to faultlessly replicate the columnar shape from the PDMS mold. Then, a new layer of the same solution was deposited under the same conditions for printing. Imprinting of sol–gel films with a PDMS mold involves the following steps. First, molds were degassed under vacuum (10 mbar) for 20 min before direct application on the as-prepared xerogel films kept in a controlled environment, without additional pressure. After imprinting, the samples were transferred to a 70 °C stove for 1 min and then to a 140 °C stove for 2 min to consolidate the xerogel films before peeling off the PDMS mold. Next, the sol–gel replicas were annealed at 430 °C for 10 min for consolidation.

### Structural characterization of nanostructured silica pillars

The microstructure of the nanostructured substrates was investigated with a FEG-SEM model Su-70 Hitachi, equipped with an EDX detector X-max 50 mm^2^ from Oxford instruments. The topography of silica pillars was studied by tapping Atomic Force Microscopy (AFM) images obtained using Park Systems NX-10 Scanning Probe Microscopy (SPM) unit.

### Cell lines and constructs

HeLa cells (a gift from B. Goud, Institut Curie, CNRS UMR 144, Paris, France) were cultured in DMEM GlutaMAX supplemented with 10% fetal calf serum and 100 U·mL^-1^ of penicillin and streptomycin at 37ºC in 5%CO_2_. Genome edited SUM159 AP-2 σ2-EGFP +/+ cells previously described^36,48^ and cultured in DMEM/F12 (1:1) GlutaMAX supplemented with 5% fetal calf serum, 100 U·mL^-1^ of penicillin and streptomycin, 1μg·mL^-1^ Hydrocortisone, 5μg·mL^-1^ insulin and 10mM HEPES at 37ºC in 5%CO_2._ All cell lines were tested negative for mycoplasma.

mCherry-MyrPalm was a gift from Gaelle Boncompain, Insitut Curie, CNRS UMR 144, Paris, France). EGFR-GFP was a gift from Alexander Sorkin (Addgene plasmid #32751). Plasmids were transfected 24h after cell seeding using JetPEI^®^ transfection reagent (Polyplus transfection^®^) according to the manufacturer’s instructions.

### Antibodies and drugs

Mouse monoclonal anti-adaptin α (Cat. Nr. 610501) was obtained from BD Transduction Laboratories (Becton Dickinson France SAS, France). Polyclonal rabbit anti-clathrin heavy chain (Cat. Nr. SAB1410212) and rat monoclonal anti-β1-integrin (Cat. Nr. MAB1997) were from Sigma. Rabbit monoclonal anti-caveolin-1 (Cat. Nr. SC-53564) was from Santa Cruz Biotechnology, Inc. Phalloidin-Atto 390 (Cat. Nr. 50556) was obtained from Sigma. Secondary antibodies were from Jackson ImmunoResearch Laboratories Inc.: Donkey anti-Mouse IgG (H+L) conjugated to Cy3 (Cat. Nr. 715-165-150) and Alexa Fluor^®^ 488 (Cat. Nr. 715-545- 150). Donkey anti-Rabbit IgG (H+L) conjugated to Cy3 (Cat. Nr. 715-165-152). Cytochalasin D (Cat. Nr. C8273), Jasplakinolide (Cat. Nr. J4580), Poly-L-lysine (Cat. Nr. P4707), Collagen, Type I solution from rat tail (Cat. Nr. C3867) and fibronectin (Cat. Nr. F1141) were purchased from Sigma-Aldrich.

### Phosphoinositide’s probes

Recombinant GST-tagged PH-domain of PLCδ detecting the membrane lipid PI(4,5)P2 was produced and conjugated to of amine-reactive Alexa Fluor 647 carboxylic acid succinimidylester (Invitrogen) as previously described^49^. Recombinant eGFP-GST-tagged PH-domain of Grp1 detecting PI(3,4,5)P3 was expressed overnight at 18 ºC using 1 mM IPTG in Escherichia coli strain BL21(DE3) and purified by affinity chromatography using glutathione Sepharose 4B beads according to the manufacturer’s instructions (GE Healthcare) in 50mM Tris at pH 8.0, 100mM NaCl. Finally, the motif was dialyzed overnight in a Slide-A-Lyzer dialysis cassette (MWCO 10,000) and the concentration was measured using a Bradford assay (Biorad).

### Immunofluorescence

Cells seeded on SiO_2_ nanostructured substrates were fixed and permeabilized in 3.7% PFA, 0.05% Triton X-100 in PBS for 20 min at room temperature, then rinsed in PBS twice and incubated for 30 min at room temperature in 1% BSA. Coverslips were then stained for the primary antibodies for 45 min. Then, secondary antibodies were incubated for 45 min. Phalloidin and PI-probes were also incubated for 45 min at a final concentration of 0.4μM and 40 μg. mL^-1^, respectively. Finally, coverslips were soaked in PBS, then in sterile water and mounted with a Mowiol^®^ 4-88 mounting medium (Polysciences, Inc.). Montage was allowed to solidify in the dark for 48h before microscope acquisitions.

### Airyscan microscopy

Images were acquired on a Zeiss LSM880 Airyscan confocal microscope (MRI facility, Montpellier). Excitations sources used were: 405 nm diode laser, an Argon laser for 488 nm and 514 nm and a Helium/Neon laser for 633 nm. Acquisitions were performed on a 63x/1.4 objective. Multidimensional acquisitions were acquired via an Airyscan detector (32-channel GaAsP photomultiplier tube (PMT) array detector). 3D images were acquired by fixing a 0.15μm z-step to generate a z-stack of images (z-stack ∼ 10) that cover the entire nanopillars height.

### Spinning disk microscopy of living cells

Live cell imaging of genome edited SUM159 AP-2-σ2-EGFP cells seeded on arrays of 1D nanopillars with *D* ∼ 600 nm was performed using a spinning disk confocal microscopy (Nikon TI Andor CSU-X1 equipped with a Yokogawa spinning disk unit (Andor)) (MRI facility, Montpellier) equipped with a 488 laser beam (60 mW). Acquisitions were performed with a 60x/1.4 objective. During imaging, cells were maintained at 37°C, 5%CO2 in an onstage incubator (Okolab). Movies were recorded with a mono dichroic mirror (488nm) and a GFP emission filter (525-30nm). Samples were exposed to laser light for 100ms every 2s for 5 min and images were acquired using an EMCCD iXon897 (Andor) camera. A 0.15μm z-step was used to cover all the nanopillars length in every z-stack acquisition.

### Fluorescence lifetime imaging microscopy of live cells

FLIM measurement of FliptR lifetime on living cells were performed as previously reported^23^. Briefly, HeLa cells were seeded on nanostructured SiO_2_ structures overnight as stated above. Then, the culture medium was replaced with the same medium containing 1.5 µM of the FliptR probe and incubated for 10 min. FLIM imaging was performed using a Nikon Eclipse Ti A1R microscope equipped with a Time Correlated Single-Photon Counting module from PicoQuant. Excitation was performed using a pulsed 485 nm laser (PicoQuant, LDH-D-C-485) operating at 20 MHz, and emission signal was collected through a bandpass 600/50 nm filter using a gated PMA hybrid detector and a TimeHarp 260 PICO board (PicoQuant). The duration of FLIM experiments was ∼ 15s. SymPhoTime 64 software (PicoQuant) was then used to fit the fluorescence decay data (from either full images or regions of interest) to a dual exponential model after deconvolution for the instrument response function.

### Fluorescence Recovery After Photobleaching of live cells

FRAP measurements where performed on a Zeiss LSM880 Airyscan confocal microscope equipped with a live cell chamber (set at 37°C and 5% CO_2_) (MRI facility, Montpellier). Excitation source was an Argon laser for 514 nm. Acquisitions were performed on a 63x/1.4 objective. Briefly, mCherry-MyrPalm labeled membranes on HeLa cells were photobleached over a rectangular area comprising ≥ 6 vertical nanopillars using 20 iterations of the 514 nm laser with 100% laser power transmission. We recorded *n* =10 pre-bleaching images and then the selected region of interest at the base of nanopillars was beached (corresponding to one z-stack). The recovery of fluorescence was traced for ≥ 25 s. After 2-5 minutes, we moved to the z plane corresponding to top of pillars and repeated the same operation (as shown in Figure S4). This routine was alternated so that, on other cells we first bleached the top of the pillars and then the base to prevent any contribution resulting from the plane that was first bleached. Fluorescence recovery and T_half_, which stands for the time point where the fluorescence is recovered to half of the maximum recovery, were calculated using the ZEN software (Zeiss), where normalized FRAP curves were fitted to model the free diffusion of one diffusing component. Data was corrected for background and incidental bleaching from imaging. Each FRAP condition reports N ≥ 5 cells.

### Data analysis and processing

The preferential localization of each cellular component of interest at the plasma membrane interface with the vertical nanopillars was determined by estimating an enrichment (*E*), as detailed as follows,

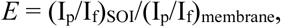

which reports the ratio between a signal of interest (SOI), at the plasma membrane bent by the pillars (Ip) versus the SOI at the flat membrane surface (If, not on pillar). This ratio was then normalized by an unspecific plasma membrane marker (e.g. mCherry-MyrPalm) on pillars and not on pillars to correct for topographical artifacts on our estimations.

The enrichment (*E*) was obtained from maximum intensity projection of z-stack images acquired by Airyscan imaging. Each acquisition was treated by using a semi-automatic detection of each SOI on pillars and not on pillars using built-in macro functions and the ImageJ software^50^.

The automated image analysis procedure calculates statistics about the average maximal height of the signal in each channel of interest. A supplementary reference channel defines the positions of the columns and their heights. A background correction is applied to each channel. Regions, that are clearly background for all z-positions, are detected. The mean intensity per z-position in the background regions is measured. A 3rd degree polynomial is fit to the data, in order to smooth it. The resulting mean background intensity is subtracted for each z-position.

An intensity threshold is calculated for each channel and a new 2D-image is created. In the new image the value of each pixel represents the maximum z-position of the voxel, for which the intensity was above the threshold. The height histograms are calculated. Height values of zero and height values at positions for which the height in the reference channel is not maximal are excluded from the calculation of the histograms. The mean height, standard-deviation and quartiles are calculated from the height-histograms. The values are reported relative to the maximum height of the reference channel. The analysis has been implemented as an ImageJ^50^ macro toolset^51^.

Two-dimensional (2D) autocorrelation analysis of maximum intensity projection of z-stack images acquired by Airyscan imaging was performed using Gwyddion (http://gwyddion.net/), a modular program for data visualization and analysis. For *N* × *N* equidistant pixels, the 2D height–height correlation function can be defined as *C*(*r*) = [*z*(*r*_0_+*r*) –⟨*z*⟩][*z*(*r*_0_) –⟨*z*⟩], where ⟨ … ⟩ means the average over all possible pairs in the matrix that are separated by a vector r = *xe*_x_ + *ye*_y_. The *z* values that are taken into account in the equation are the deviation from the average height ⟨*z*⟩52.

### Data representation and statistical analysis

3D representations of Airyscan images were generated with the 3/4D visualization and analysis software Imaris (Oxford Instruments).

Lifetime plots were generated using web-based data visualization tools (Shiny apps, https://huygens.science.uva.nl): PlotsOfData^53^. Enrichment plots were generated using Prism GraphPad software. Data is represented as dot plots reflecting the data distribution and the median of the data. Vertical bars indicate for each median the 95% confidence interval.

Statistical analysis was performed using Prism GraphPad software. Statistical test, significance and data points (*n*) are specified on each graphical representation. Experiments represent *N* ≥ 3 replicates.

## Supporting information

Contents additional data about the fabrication of nanostructured substrates and characterization of samples.

## Acknowledgements

The authors thank D. Montero for performing the FEGSEM images. B. Charlot and R. Desgarceaux from IES for providing the silicon mold. C. Cazevielle (MRI-COMET, Montpellier) for assistance with SEM images. C. Favard, D. Muriaux, and N. Sauvonnet for scientific discussions. We acknowledge the imaging facility MRI, member of the national infrastructure France-BioImaging infrastructure supported by the French National Research Agency (ANR-10-INBS-04, "Investments for the future"). The authors are grateful to A. Roux for the support provide to this work. L.P. acknowledges the ATIP-Avenir program for financial support. A.C-G. acknowledges the financial support from the European Research Council (ERC) under the European Union’s Horizon 2020 research and innovation program (No.803004) and the French Agence Nationale de la Recherche (ANR), project Q-NOSS ANR ANR-16-CE09-0006-01; The FEGSEM instrumentation was facilitated by the Institut des Matériaux de Paris Centre (IMPC FR2482) and was funded by Sorbonne Université, CNRS and by the C’Nano projects of the Région Ile-de-France.

## Author contributions

T.S., D.S, R.R., A.C-D. performed experiments and analyzed results. F.E-A. and J.V. generated the PI-probes. S.R. and V.B designed the semi-automated and automated analysis of images. R.G. generated the genome-edited SUM-159 cell lines. M.M. and S.M. synthetized the FliptR probe. A.C-G. and L.P. supervised the study, designed experiments, performed experiments and wrote the manuscript. All authors contributed to the final version of the paper.

## TOC figure

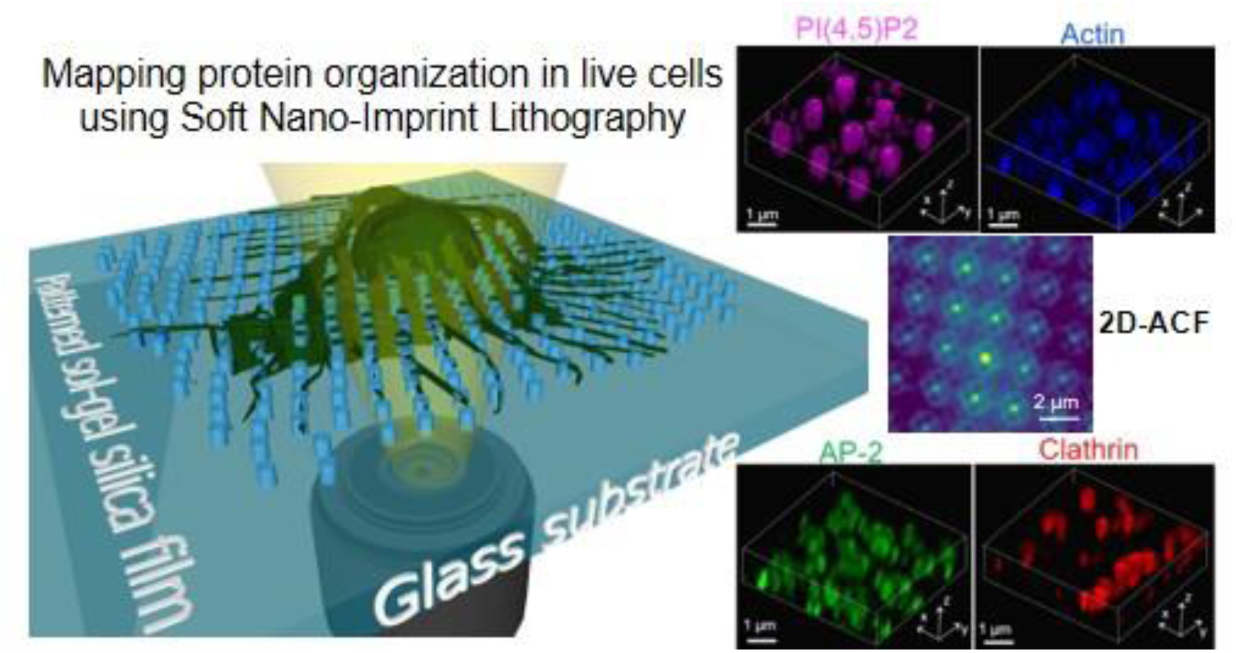

